# Audiotactile stimulation can improve syllable discrimination through multisensory integration in the theta frequency band

**DOI:** 10.1101/2023.05.31.543034

**Authors:** Pierre Guilleminot, Cosima Graef, Emilia Butters, Tobias Reichenbach

## Abstract

Syllables are an essential building block of speech. We recently showed that tactile stimuli linked to the perceptual centers of syllables in continuous speech can improve speech comprehension. The rate of syllables lies in the theta frequency range, between 4 and 8 Hz, and the behavioural effect appears linked to multisensory integration in this frequency band. Because this neural activity may be oscillatory, we hypothesized that a behavioural effect may also occur not only while but also after this activity has been evoked or entrained through vibrotactile pulses. Here we show that audiotactile integration regarding the perception of single syllables, both on the neural and on the behavioural level, is consistent with this hypothesis. We first stimulated subjects with a series of vibrotactile pulses and then presented them with a syllable in background noise. We show that, at a delay of 200 ms after the last vibrotactile pulse, audiotactile integration still occurred in the theta band and syllable discrimination was enhanced. Moreover, the dependence of both the neural multisensory integration as well as of the behavioral discrimination on the delay of the audio signal with respect to the last tactile pulse was consistent with a damped oscillation. In addition, the multisensory gain is correlated with the syllable discrimination score. Our results therefore evidence the role of the theta band in audiotactile integration and provide evidence that these effect may involve oscillatory activity that still persists after the tactile stimulation.

## Introduction

Speech is built from units of increasing duration and complexity, from phonemes to syllables, words and sentences. Extraction of the linguistic information in the brain and its subsequent processing requires some segmentation of the acoustic signal into these units [1, 2]. This segmentation presumably relies on cortical activity that tracks the rhythms set by the different speech units [3].

The theta frequency range of cortical activity, between 4 Hz and 8 Hz, appears especially important as it corresponds to the rhythm of syllables and has been demonstrated to reflect several speech-related processes [4, 5, 6]. Moreover, influencing the cortical tracking in the theta frequency range through transcranial alternating current stimulation (tACS) has been found to modulate speech comprehension [7]. The delta frequency range, between 1 Hz and 4 Hz, is important as well as it corresponds to the rhythm of words and has been linked to syntactic and semantic information [8, 9]. However, it should be noted that influencing the cortical tracking in that range using tACS did not modulate speech comprehension [7].

The neural activity in the theta band is also thought to support multisensory integration for speech comprehension as well as for attention to speech [10, 11]. Because the cortical activity in frequency bands such as the theta band is often assumed to be oscillatory, the multisensory integration has been hyphothesized to work based on a phase reset mechanism. Under this hypothesis, the ongoing oscillations in the theta frequency band are reset through a stimulus, yielding and enhancing the observed cortical tracking of amplitude changes in a continuous stimulus such as speech [12, 13]. This hypothesis is supported by the observation of resets of activity in the auditory cortex through visual inputs [14]. Regarding speech processing, the neural activity linked to speech rhythms may accordingly be reset by stimuli in other sensory modalities, possibly impeding or enhancing speech comprehension [15, 16]. In addition to this bottom-up mechanism, multisensory effects may also involve top-down attentional modulation, which may involve alpha band activity [17].

Tactile stimuli have been found to elicit activity in the auditory cortex, and this neural activity becomes integrated with auditory activities at similar latencies as the processing of the auditory information itself [18, 19, 20, 21]. Somatosensory stimuli, in particular, have been shown to reset the phase of ongoing oscillatory activity in the auditory cortex [22] as well as to modulate the cortical tracking of the amplitude fluctuations in speech [23].

Regarding behavior, studies have found that speech comprehension can be modulated using tactile stimuli derived from slow rhythms of speech, generally after training of the subjects [24, 25]. We showed recently that sparse vibrotactile signals delivered at the rhythm of syllables can modulate both speech comprehension and certain neural responses to speech [26]. In particular, the tactile stimuli can both enhance as well as reduce speech comprehension, depending on the delay between the vibrotactile pulses and the perceptual centers of the syllables. The dependence of the speech comprehension score on the delay follows a sinusoidal function with a frequency that lies in the theta range, and speech comprehension is maximal when the oscillations reach their peak. This behaviour might reflect a multisensory phase reset mechanism in which oscillatory neural activity in the auditory cortex could be reset in phase through the tactile stimuli. The tactile stimulation might thereby aid the segmentation of the neural activity into different syllables and thus support syllable parsing. Alternatively, it might lead to the syllables occurring at phases of the neural activity that provide the highest sensitivity in neural processing, and thus aid the processing of the information in the individual syllables. However, since this study employed continuous speech, it remains unclear whether the observed sinusoidal dependence reflects oscillatory brain activity or rather a regularity in the timing of the syllables [27].

Here, we aim to further investigate the audiotactile integration. In particular, we wish to test whether tactile stimuli can affect the comprehension of a single syllable even after the tactile stimuli have ended, as would be expected if the stimuli entrained ongoing oscillatory brain activity. To do so, we restrict speech comprehension to a syllable discrimination task. Indeed, by stimulating the activity in the auditory cortex through vibrotactile stimulation at a naturally occurring syllabic rate, we might observe an improvement in syllable discrimination if the audiotactile speech enhancement worked through setting the neural activity in the auditory cortex to occur at a phase of maximal sensitivity with respect to the auditory signal. We therefore aim to investigate both behaviour, through measuring the ability of subjects to discriminate between two similarly-sounding syllables, as well as electrophysiology, to elucidate neural correlates of audiotactile integration.

## Materials and Methods

### Participants

Sixteen subjects (aged 22 *±* 2 years old, seven of them female) participated in the experiment. The number of participants was chosen based on previous related work [15, 7, 24, 26]. All subjects were native English speakers, right-handed, and had no history of neurological disorders or hearing impairment. No prior training regarding the tactile stimulation, with or without audio, was given beforehand. Because of equipment failure, one of the subjects’ EEG could not be recorded. Volunteers gave their written informed consent prior to the experiment. The research was approved by the Imperial College Research Ethics Committee (ICREC) with the reference: 19IC5388 NoA 1.

### Experimental Design

Subjects were first stimulated with five subsequent vibrotactile pulses that occurred at a rate of 5 Hz, that is, two successive pulses were spaced 200 ms apart (Figure 1). Background noise was presented continuously throughout the trial. After a delay of 150 ms to 300 ms from the time of the last pulses, subjects heard a short syllable. They then had to choose which of two similarly-sounding syllables was presented, a task for which they were allowed to take as much time as needed.

**Figure 1:**
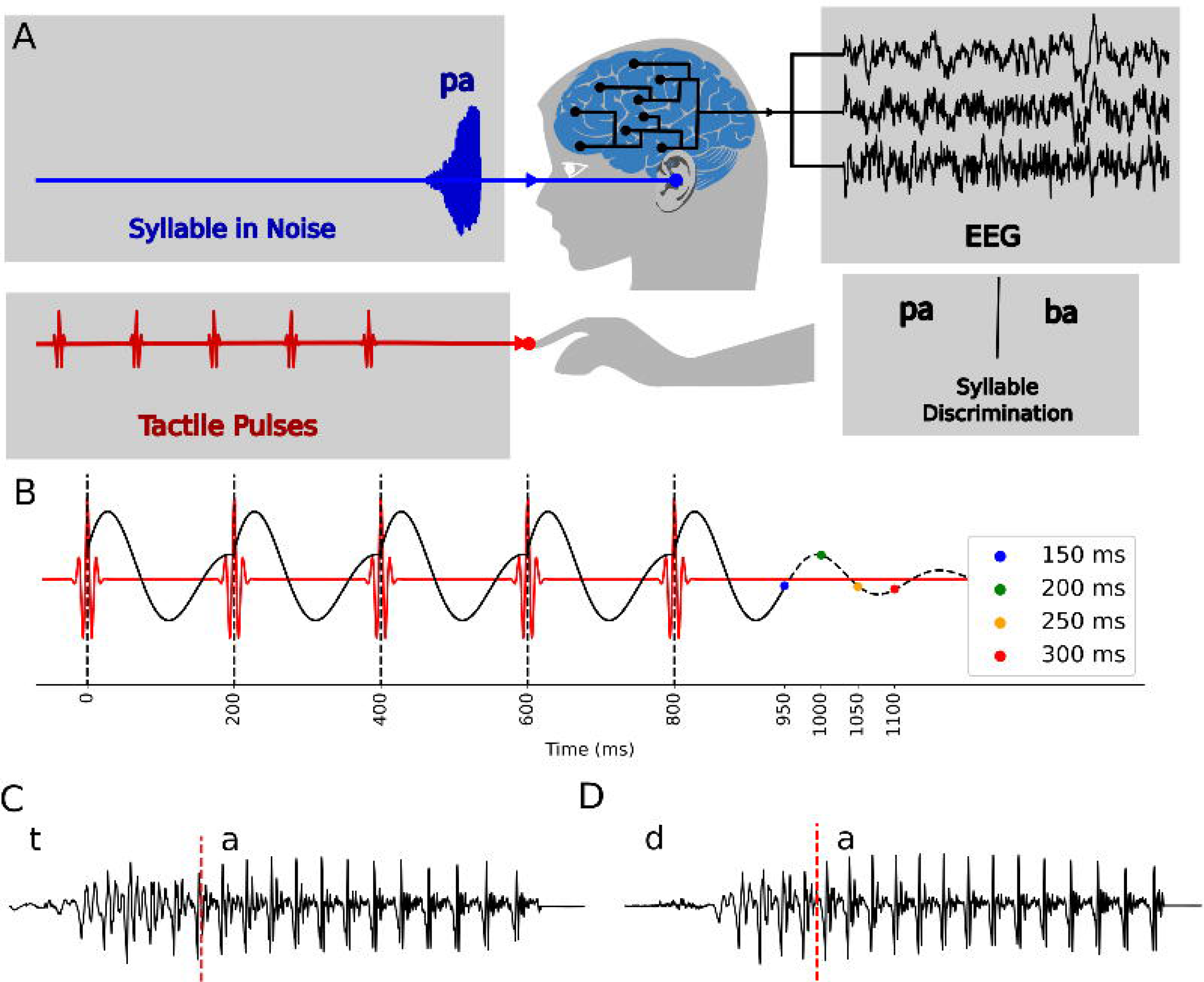
Experimental setup. (**A**) Each subject was first stimulated with vibrotactile pulses at their right index finger, and subsequently heard a syllable in noise that they had to discriminate from another similarly-sounding syllable. During the experiment, brain activity was recorded using EEG. (**B**) The stimuli consisted of five consecutive vibrotactile pulses (red line) at a rate of 5 Hz, followed by a syllable whose onset occurred 150 ms, 200 ms, 250 ms, or 300 ms (blue, green, yellow and red disk) after the last vibrotactile pulse. Putative oscillatory neural activity in the theta range might be reset by the vibrotactile pulses and decay after the last pulse (black line). (**C,D**) The waveforms for one pair of syllables: ta and da. The red line marks 50ms.

Electroencephalographic (EEG) responses were recorded throughout the experiment, starting before the delivery of the first vibrotactile pulse and ending after the presentation of the syllable.

### Data and Code availability

The data is accessible from a public database (https://zenodo.org/record/7544913) [28] and code is accessible on github (https://github.com/phg17/ATacSy).

### Hardware

All signals were created digitally on a PC (Windows 7 operating system). They were then synchronized and converted into analogue signals through the RX8 multi-I/O processor device (Tucker-Davis Technologies, U.S.A.) at the sampling rate of 39,062.5 Hz. Subjects listened to the acoustic stimuli through insert earphones (ER-2, Etymotic Research, U.S.A.) placed in their ear canals. For tactile stimulation, subjects were holding a vibrotactile motor (Tactuator MM3C-HF, TactileLabs, Canada) between the thumb and index finger of their right hand.

EEG signals were acquired using 64 active electrodes (actiCAP, BrainProducts, Germany) and a multichannel EEG amplifier (actiCHamp, BrainProducts, Germany). For a precise synchronization of the sensory stimuli and the EEG response, the delivered signals were recorded together with the EEG signals (StimTrak, BrainProducts, Germany), and each trial was preceeded by a trigger signal sent to the TDT. The tactile and audio signals were sent using the multi-I/O-processor TDT RX8 (Tucker-Davis Technologies, USA), thus ensuring their synchronization with each other.

### Acoustic Stimuli

In each trial of the experiment, subjects first experienced five vibrotactile pulses, listened to a syllable in noise, and then selected the presented syllable from two possible choices. The two syllables from which the subject could choose in a given trial were displayed on the left and right side of a monitor, and subjects were asked to indicate their choice by pressing the left or right key respectively on the keyboard. The syllables were always presented within one of the following three pairs: (pa, ba), (ta,da), and (ga, ka). These syllables all have a similar manner of articulation, and the two syllables of a given pair also possess a similar place of articulation [29]. Indeed, all of the syllables are constructed from a stop consonant and the only difference between two members of a pair is whether they are voiced or unvoiced. Each member of the pairs can therefore be easily confused for the other, making them good candidates for a discrimination task.

The syllables were converted to both an artificial female and a male voice using the text-to-speech software TextAloud at a sampling rate of 44,100 Hz. The audio stimuli were manipulated using Praat to have the same duration and intensity [30]. The syllables thus created were modified so that the transition of consonant (’d’, ’t’, ’p’, ’b’, ’g, ’k’) to vowel (always ’a’) happened 50 ms after the start of the syllable, i.e. 50 ms after the first change in amplitude (Fig.2C,D). The information that allowed for syllable discrimination was therefore contained within the first 50 ms. This duration was chosen because of the audiotactile lags defined below that are separated by 50 ms each. Because the information on syllable discrimination was contained in a 50 ms time window, sampling the lags between tactile stimuli and the audio stimuli at this temporal spacing allowed to explore meaningful delays between these two types of stimuli.

**Figure 2:**
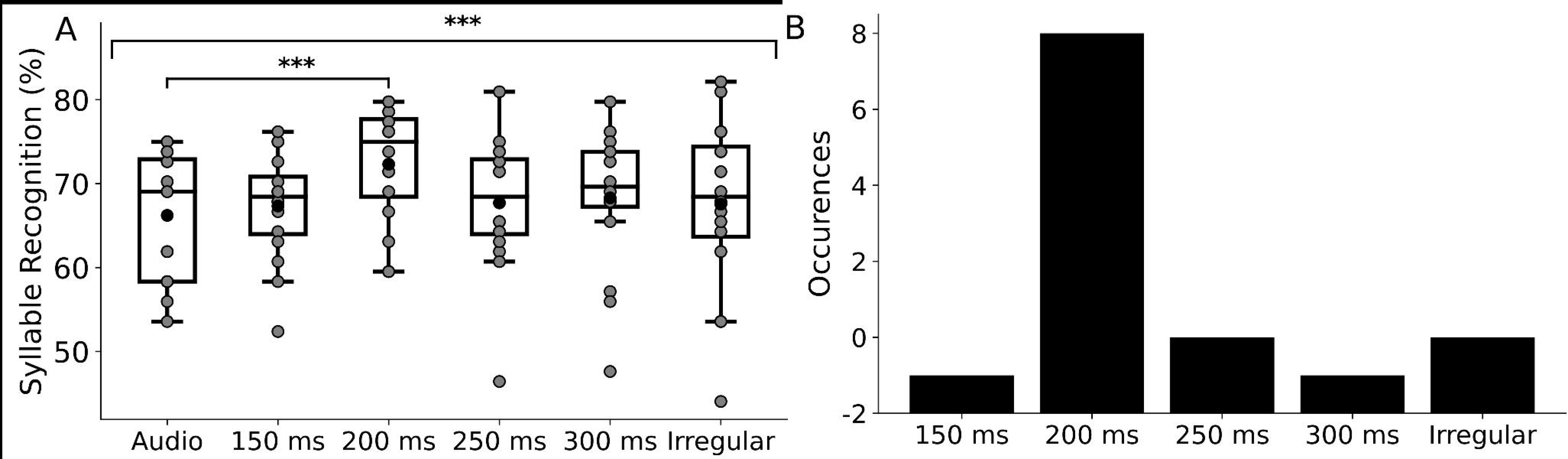
Syllable recognition scores for the different conditions. (**A**) The scores differed significantly between the conditions. In particular, syllable recognition at an audiotactile lag of 200 ms was significantly higher than in the audio-only condition. (**B**) A histogram of the number of occurrences at which a certain condition yielded the best syllable recognition per subject showed that the audiotactile lag of 200 ms gave, for most subjects, the highest score.

The signal was then mixed with speech-shaped noise at a signal-to-noise ratio of -3 dB. The speech-shaped noise was generated so that it covered the same spectral content as the target speech. The resulting audio signals were delivered to the subjects at an intensity of 75 dB SPL, deemed comfortable by the volunteers. We refer to this signal as the *audio stimuli* or the *syllables* in the rest of the paper.

### Tactile Stimuli

The tactile stimuli consisted of five vibrotactile pulses that were presented in succession at a frequency of 5 Hz, resulting in a duration of 800 ms between the first and the last pulse. In addition to this regular train of vibrotactile pulses, we created irregular pulse trains. The time between two successive pulses was thereby drawn from a uniform distribution ranging from 50 ms to 300 ms. The total duration of the obtained irregular pulse trains ranged from 700 to 900 ms. The onset of the syllable was delivered 1000 ms after the first vibrotactile pulse and thus between 100 ms and 300 ms after the last vibrotactile pulse.

The temporal waveform *ψ*(*t*) of the individual pulses was designed as real Morlet wavelets:

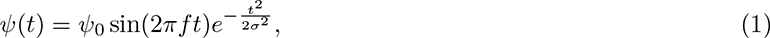

in which *t* denotes time. The amplitude was set at *ψ*_0_ = 1.4 V, the carrier frequency *f* at 80 Hz and the width *σ* at 7.5 ms. We refer to this signal as the *tactile stimuli* or the *vibrotactile pulses* in the rest of the paper.

### Delays of the sound signal and control conditions

When presenting the syllable, we considered different time shifts between the latest vibrotactile pulse and the onset of the syllable, which we refer to as the *audiotactile lags*. We expected that the audiotactile integration would involve neural activity in the theta frequency range, in which the periods of oscillation range from 125 ms to 250 ms. Investigating a putative oscillatory component in the neural, and respectively in the behavioural responses to the syllables, therefore, required delays that would lie in the second period of the oscillations following the last vibrotactile pulse, but not later so that the oscillatory activity had not yet dissipated.

We accordingly chose lags of 150 ms, 200 ms, 250 ms, and 300 ms after the last vibrotactile pulse (Figure 1). This resulted in four audiotactile conditions. A fifth condition was obtained by starting with an irregular train of vibrotactile pulses followed by the presentation of a syllable that occurred randomly between 100 ms and 300 ms after the last pulse. Whether regular or not, conditions in which both tactile pulses and syllables are presented are referred to as *audiotactile conditions*. Finally, as a control, we used an audio-only condition where no vibrotactile pulses were presented. We therefore obtained a total of six different conditions, one of which was irregular, in order to study if the simple addition of tactile stimulation, with no set rhythm, could elicit an effect. This was motivated by previous results that demonstrated a supra-additive effect on envelope tracking in the presence of mismatched tactile stimuli [26].

### Data collection

The data collection was carried out in a single session for each participant. In each trial the subjects were presented with a single syllable in one of the six conditions, that is, with prior tactile stimulation except in the audio-only condition. Before each trial, a window of text notified subjects of whether the condition was going to be an audiotactile or the audio-only condition so they would not be surprised by the presence or absence of the vibrotactile pulses. This was done in order to avoid any effects of the subjects’ surprise on the EEG signal [31]. Moreover, during a pilot study in which subject were not informed about the nature of the forthcoming stimulus, subjects tended to interpret the absence of tactile stimuli during the audio-only condition as a dysfunction, impeding their focus.

Each trial, audio or audiotactile, started with the presentation of speech-shaped noise for a random duration between 500 ms and 1 s. This was done to get the subjects used to the intensity of the audio stimulus at each trial, as well as randomizing the time at which the syllables were presented after the start of the trial. Using a random duration avoided subjects using the start of the noise presentation to infer the exact timing of the upcoming syllable. In the audiotactile conditions, the vibrotactile pulses were then sent, followed by the syllable. In the case of the audio-only condition, no vibrotactile pulses were presented but noise was played for the same duration. Finally, after the syllable had been played, there was a 500 ms pause before the choice of two syllables was displayed on the monitor. Prior to this choice, the monitor continuously displayed a fixation cross that subjects were asked to focus on.

Each unique combination of syllable (pa, ba, ka, ga, ta, da), condition (1 audio-only, 5 audiotactile), and voice (female, male) was presented seven times, resulting in a total of 504 trials. During the whole duration of the experiment, we recorded the subject’s EEG.

We did not measure the participants’ reaction time since we deliberately did not want to stress them regarding fast responses. In fact, we informed the participants that they could take as much time as needed to provide a response.

### Preprocessing of EEG recordings

We recorded the EEG data at a sampling rate of 1,000 Hz using 64 electrodes in an extended 10-20 system referenced to the vertex (Cz). During the preparation of the EEG recordings for a particular subject, the maximal impedance was kept under 10 *k*Ω. The EEG data was then band-pass filtered between 0.1 Hz and 32 Hz (one- pass, zero-phase, non-causal FIR bandpass filter of order 33,000). For time-frequency analysis, the EEG data was resampled at 100 Hz.

### Statistical Testing of the Behavioural Results

For the behavioral results on syllable discrimination, we first assessed if the results differed significantly between the six experimental conditions through a Friedman test. If an effect was detected, we then used multiple post- hoc Wilcoxon tests between the audio-only condition that served as a control condition and all of the audiotactile conditions. The resulting *p*-values were corrected for multiple comparisons using the conservative Bonferroni correc- tion [32]. We also computed the condition at which the maximum score was obtained for each subject. We then tested whether the obtained distribution could be obtained via a uniform distribution using a chi-squared test.

### Statistical Tests for the EEG Data

In order to test an effect of the tactile stimulation on the neural response to the syllable, we investigated the EEG signal after the syllable had been presented. In particular, we chose a time window between 50 ms and 550 ms after the onset of the syllable. Somatosensory evoked potentials range from 0 ms to 200 ms post-stimulus [33, 34]. Therefore this choice of window allows to avoid late-evoked activity contamination from the tactile stimuli while still studying the entire neural response to the syllable.

In this time window, we studied the EEG power in different scalp areas and in different frequency bands. In particular, we focused on the power in the delta frequency band (1 *−* 4 Hz) and in the theta frequency band (4 *−* 8 Hz) in the lateral temporal areas as they have been found to play important roles in speech processing [4, 35, 36]. The theta band is especially relevant due to its role found in audiotactile integration of continuous speech [26]. In addition, we considered the power in the alpha frequency band (8 *−* 12 Hz) in the frontal and central areas, as it may reflect top-down multisensory processes i.e. attentional modulation [17].

The EEG signals in these scalp areas and frequency bands were compared between the audio-only control condition and each of the five audiotactile conditions. The comparison was obtained using a threshold-free cluster enhancement method using the Morlet transform [37, 38]. For each combination of the five audiotactile conditions and the six space-frequency foci of interest, the clustering method yielded a collection of *p*-values that were already corrected for multiple comparisons. We selected the minimal *p*-value for each of these 30 combination, and then corrected for multiple comparisons across the 30 combinations through the Bonferroni correction [32].

Although the threshold-free cluster enhancement method allows for interpretability of the obtained results, it is limited in terms of resolution and conclusions about the exact extent of latency, location, or frequency band [39, 40,

41]. However, the cluster enhancement method can reliably identify the frequencies at which the neural responses in the audiotactile condition and in the audio-only condition differ the most.

### Damped Oscillator Model

We hypothesized that the tactile stimulation can reset the phase of intrinsic neural oscillations. After the last vibrotactile pulse, we assumed that the intrinsic oscillations would slowly dissipate, following damped oscillations (Fig.1B). We therefore modeled both the neural and the behavioral data obtained in the audiotactile conditions with different lags through damped oscillations:

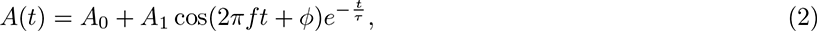

in which *t* denotes the audiotactile lag. In order to reduce subject-specific differences, the data points were scaled so that they would have a standard deviation of 1 and be centered around 0 for each subject, such that *A*_0_ = 0. *A*_1_ corresponds to the amplitude of the oscillation, while *f* represents the oscillation frequency and *ϕ* a phase shift.

Finally, *τ* is the decay time of the oscillation.

This simple model is represented in Fig. 1B, where an oscillation is reset by tactile stimulation. The oscillation at frequency *f* may be viewed as capturing some aspects of neural activity in the auditory cortex. After the last stimulation the oscillation is assumed to decay in time due to dissipation. Our experiment assumes that the decay time *τ* is long enough that some oscillatory activity can be observed past a single period. Finally, the phase *ϕ* denotes the phase of the oscillations after the reset. For example, Fig. 1B represents a low negative phase 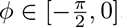. The neural activity shortly afterwards reaches a maximum which may mean optimal sensitivity to acoustic processing.

For the frequency *f*, we considered the values that the cluster enhancement method identified as being most informative about the differences in the EEG signals between the audiotactile conditions and the audio-only condition. We made this choice since our analysis showed these frequencies to be related to audiotactile integration in the brain.

We obtained 64 data points (4 audiotactile lags for 16 subjects) for the behavioural measure and 60 data points (4 audiotactile lags for 15 subjects) for the electrophysiological one. The fitting procedure was done in two steps, as per previously presented [26, 7, 42]. First, we fitted the amplitude, phase, and decay time through least-square optimization using the trust-region reflective method and a linear loss. The parameters were fitted with the following bounds: *A*_1_ *∈* [0, 100], *ϕ ∈* [*−π, π*], *τ ∈* [0, 1000] ms. We later used the values of *ϕ* and *τ* found this way. Second, the statistical significance of the obtained fit was assessed through Hubert robust linear regression, as it accounts for possible outliers, using *A*_1_ as the regression variable and a significance criterion of *α* = 0.05 [43].

### Correlation between Electrophysiology and Behavior

Finally, we wanted to relate the behavioural results to the electrophysiological one. In particular, we sought to capture whether there was a common relation between the syllable recogntion and the neural audiotactile gain within individuals. To that end, we used repeated measures correlation [44]. We therefore considered each of the different audiotactile condition, including the irregular one, as a measure for each of subjects, resulting in a total of 75 data points (15 subjects x 5 audiotactile conditions).

## Results

### Behavioral Assessment

We first determined the average syllable discrimination score across all subjects and all conditions. On average, subjects recognized 70 *±* 1% of the syllables correctly (mean and standard error of the mean).

We then studied the variation of this score between the different conditions. A Friedman test revealed a significant difference between the scores obtained in the various experimental conditions (*p <* 0.0005). We therefore tested the syllable discrimination score in each audiotactile condition against the score obtained in the control audio-only control condition using post-hoc pairwise tests. These tests revealed that the syllable discrimination score in audiotactile condition with an audiotactile lag of 200 ms was significantly higher than the score in the audio-only condition (*p* = 0.0006, Bonferroni correction for multiple comparisons, Fig. 2A). On average, subjects scored 68 *±* 1% in the control condition, whereas they achieved an average score of 74*±*1% in the audiotactile condition with an audiotactile lag of 200 ms. No significant difference emerged for any of the other comparisons (*p >* 0.1).

To further analyze the difference between the scores obtained with the audiotactile lag of 200 ms and the ones in the audio-only condition, we computed the difference between these scores for each syllable separately and compared them. A Friedman test did, however, not find a significant variation in the score differences between the different syllables (*p* = 0.49).

Finally, as an additional measure of audiotactile integration, we computed the audiotactile condition for each subject at which they obtained the best score. The obtained distribution was significantly non-uniform(*p* = 0.001, Fig. 2B). The condition that yielded the highest syllable recognition score for most subjects was the one with the audiotactile lag of 200 ms.

### Electrophysiological Responses

To study the neural responses to the sensory stimuli, we first investigated the responses to the five vibrotactile pulses. We obtained five consecutive peaks, spaced 200 ms apart (Fig. 3A). Each peak occurred at a delay of 48 ms after the corresponding pulse. The topographic plots showed a response in the left hemisphere, with a dipole bipolar pattern that suggested a source in the left somatosensory cortex. This spatiotemporal pattern was confirmed in the average response across the five pulses (Fig. 3B). To assess the amplitudes of the five peaks, we computed the global field power of the EEG signal in a region of interest (FC1, FC3, F3, F1, CP5, CP3, P5, P3). (Fig. 3C).

**Figure 3:**
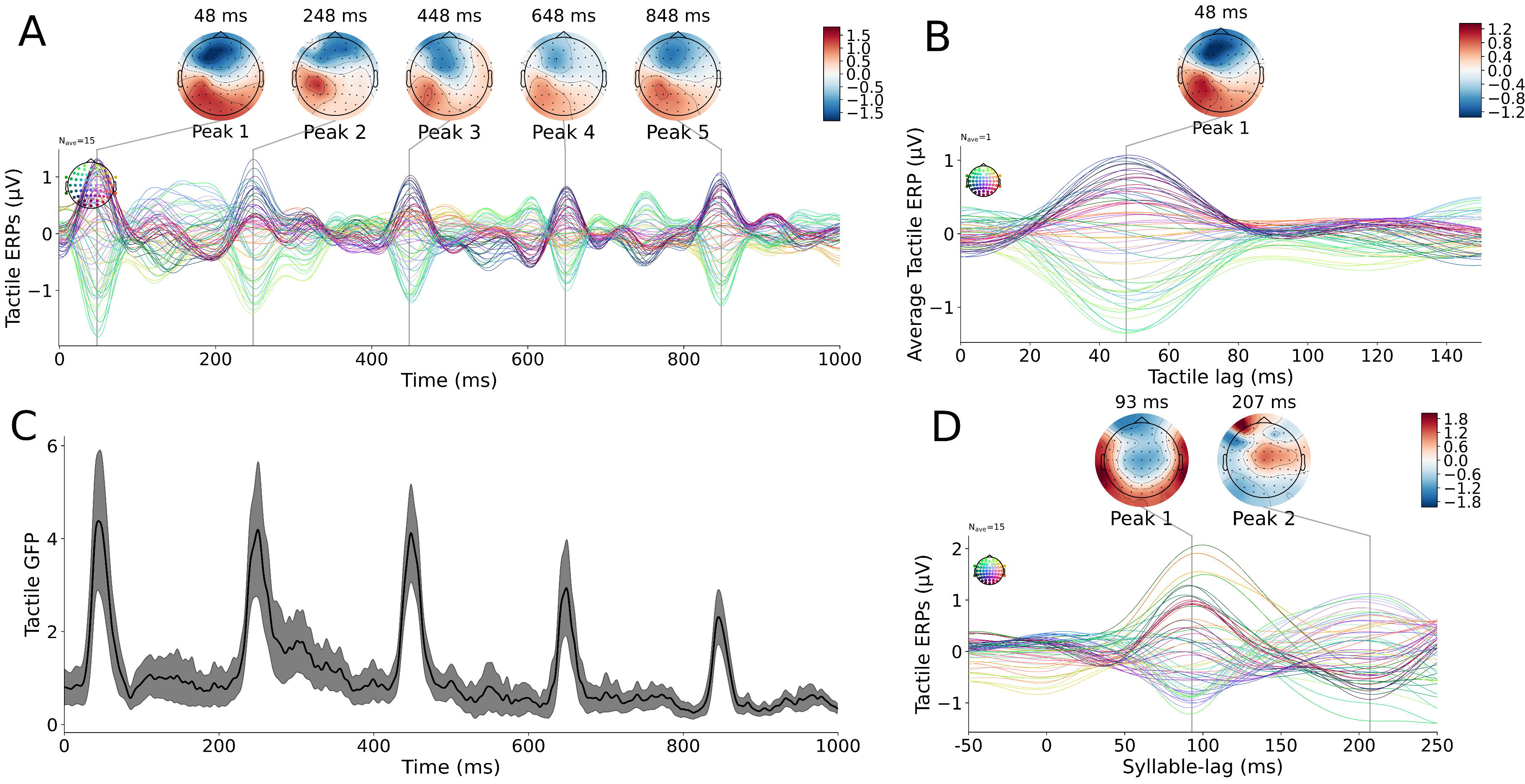
EEG responses to the audiotactile stimulation. (**A**) The EEG responses evoked by the vibrotactile pulses suggest a source in the left somatosensory cortex (average over all audiotactile conditions, time is relative to the first pulse). (**B**) The EEG signal in a response to a single vibrotactile pulse. This was obtained by averaging the individual responses to each of the five pulses together. The result showed a clear left-lateralized response in the somatosensory area. (**C**) The global field power, that is, the average power in the EEG signal across all electrodes, evidenced a decay in the amplitude between the first and the last pulse (black line, mean; black shading, standard error of the mean). (**D**) The EEG signal in response to the syllable in the audio-only condition, with time measured relative to the syllable onset, showed two peaks, a first at 93 ms and a second at 207 ms.

Finally, we assessed the EEG response to the syllable in the audio-only condition (Fig. 3D). We found two peaks, at delays of 93 ms and 207 ms after the onset of the syllable. The first peak showed a larger amplitude with a symmetric topography while the second peak displayed a left-lateralized topography.

### Audiotactile gain in the EEG response

To assess whether there was an audiotactile gain in the EEG response, we considered the EEG signal from before the onset of the syllable to a certain period afterwards. We computed the space-time-frequency response of each audiotactile condition, and subtracted the response of the audio-only condition. This yielded an estimate of the audiotactile gain provided by the prior presentation of the vibrotactile pulses.

We focused on specific regions of interest in the channel-frequency space (Fig. 4A). First, we were interested in the delta and theta frequency band in the left and right lateral temporal areas. Second, we also investigated the responses in the alpha band in the frontal and central areas.

**Figure 4:**
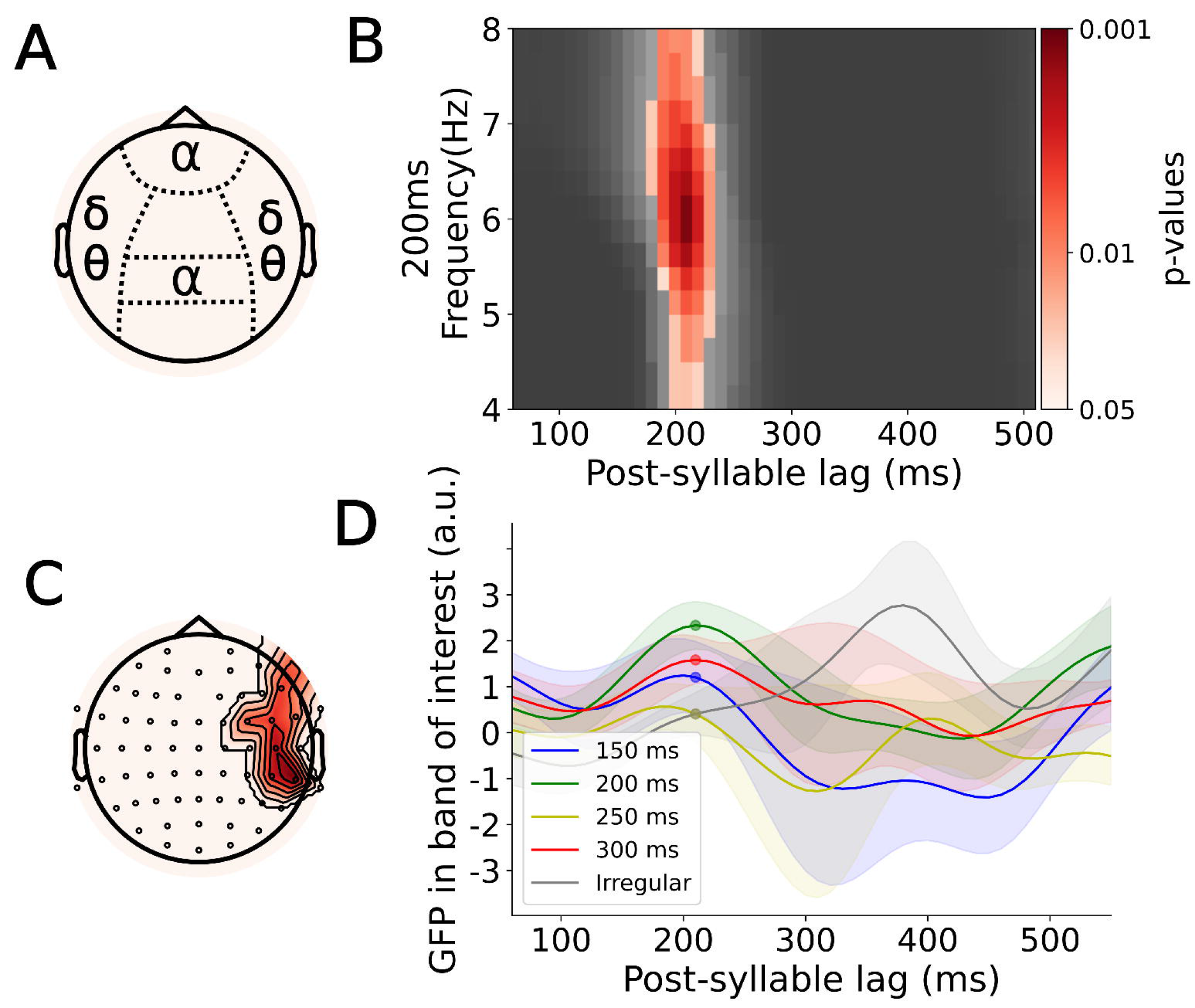
EEG markers of audiotactile integration. (**A**) A map of the different scalp areas and associated frequency bands for which we assessed potential differences in the neural response in the audiotactile conditions to that in the audio-only condition. (**B**) A threshold-free cluster enhancement method applied to EEG data in the theta band and in the right temporal area as a function of time and frequency showed a significant signal around a lag of 200 ms and around a frequency of 6 Hz. The lag was measured with respect to the onset of the syllable. Insignificant *p*-values are shown in black. (**C**) Results from the threshold-free cluster enhancement for the theta-band in the right temporal area as a function of EEG channels at the maximum given by the previous graph (**B**). (**D**) The power in the EEG signals at 6 Hz in the right temporal area as a function of the post-syllable lag for different audiotactile lags. A maximum occurred at a post-syllable lag of 200 ms. The value of the EEG power at this lag, indicated by the colored disks, was subsequently used as an electrophysiological marker of audiotactile integration.

We found only one space-time-frequency cluster at which a statistically-significant audiotactile gain emerged. This cluster was located on the right lateral area, contained frequencies in the theta range and occurred at a delay of 200 ms after the syllable onset (*p* = 0.036, Bonferroni correction for multiple comparisons, Fig. 4B,C). The cluster corresponding to this particular set of frequency (theta range), condition (audiotactile with a latency of 200 ms) and location (right lateral area) is shown in Fig. 4. No other cluster proved statistically significant after multiple test correction (*p >* 0.13).

The center frequency of the cluster at which the audiotactile gain emerged was 6 Hz. We computed the global field power at this frequency in each audiotactile condition and subtracted the corresponding power in the audio-only condition (Fig. 5D). Because the audiotactile gain occurred around 200 ms after the syllable onset, we extracted the power difference at this latency. Since this quantity represents a marker of audiotactile integration, we refer to it as the *neural audiotactile gain*.

**Figure 5:**
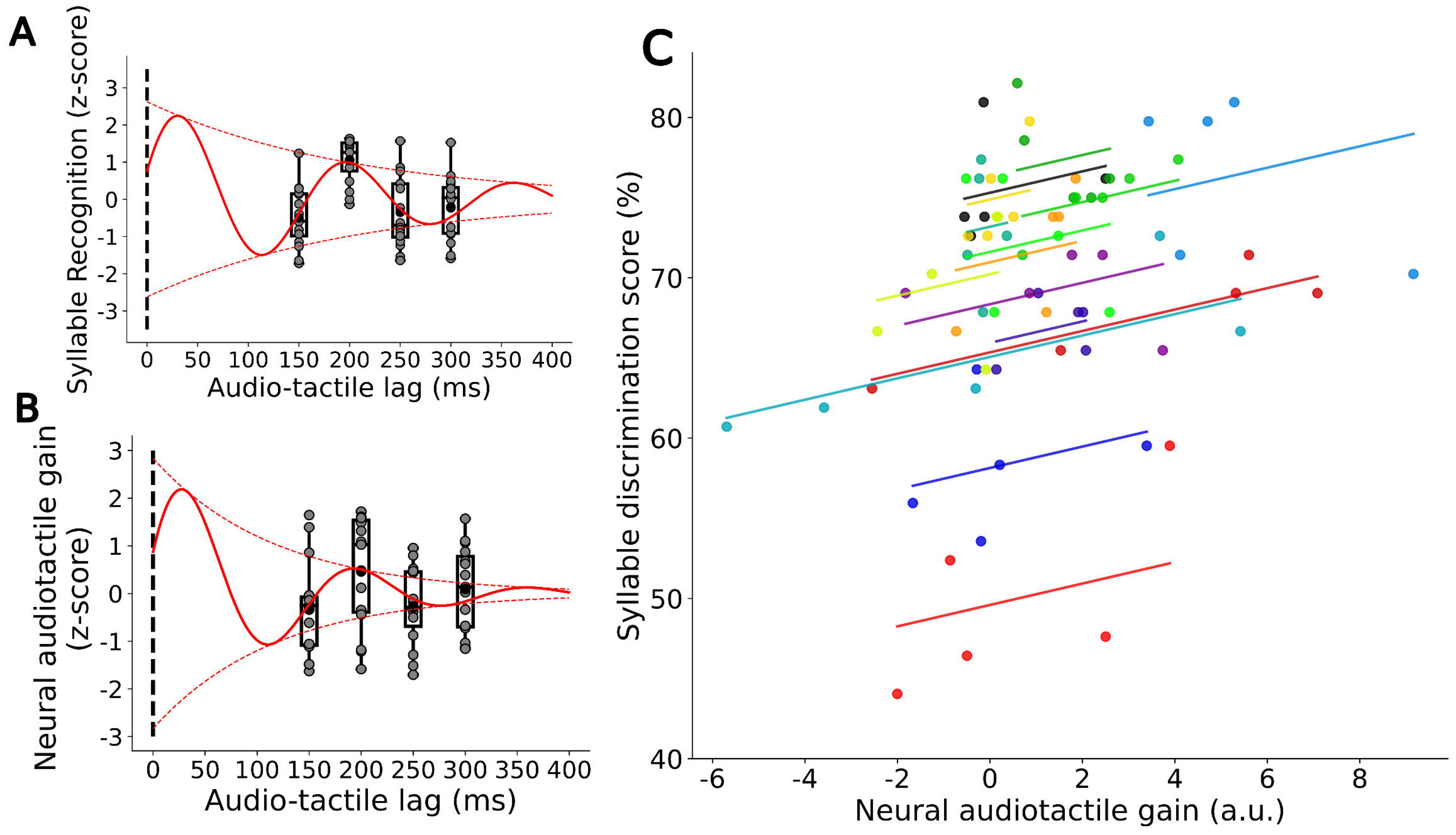
Dependence of behavioral and electrophysiological results on the audiotactile lag and on each other. (**A**) The dependence of the syllable discrimination scores (box plots, disks show results from individual subjects) on the audiotactile lag could be modeled as a damped oscillation (red line). (**B**) Similarly, the dependence of the electrophysiological marker of audiotactile integration, the neural audiotactile gain (box plots, disks show results from individual subjects), on the audiotactile lag could be fitted by a damped oscillation (red line). (**C**) The behavioral and the neural measure were correlated with each other across the different subjects and experimental conditions (red line). The average value for each audiotactile lag is shown as a colored disk.

### Time dependency of the syllable recognition score and the neural audiotactile gain

Having identified a behavioral quantity that informs on the audiotactile integration, the syllable recognition score, and an electrophysiological one, the neural audiotactile gain, we wondered how both depended on the delay at which the syllable occurred after the last vibrotactile pulse. Since we hypothesized that speech processing employs neural oscillations, the phase of which is reset by the tactile stimuli, we assumed that these oscillations would disperse and hence decay in amplitude after the last vibrotactile pulse.

We therefore described the time dependence of both the behavioural as well as of the electrophysiological quantity through a damped oscillation. We chose an oscillation at a frequency of 6 Hz, the one at which the neural response was observed, to display audiotactile gain. The other parameters of the damped oscillations were fitted to the data. We obtained a significant fit with a phase shift of *−*1.27 rad and a decay time of 166 ms (*p <* 1*e −* 6, *R*^2^ = 0.358, (Fig. 5A)). The first and second peak of the fitted curve occur at 31 ms and 197 ms after the last vibrotactile pulse, respectively.

We analogously fitted the time dependency of the neural audiotactile gain through a dampened oscillation with a frequency of 6 Hz. We obtained a phase shift of *−*1.26 rad, and a decay time of 116 ms (*p* = 0.027, *R*^2^ = 0.078, Fig. 5B). The first and second peak of the obtained curve emerged at delays of 27 ms and 193 ms after the last pulse, very similar to those obtained for the syllable recognition score.

### Correlation between Behavior and Electrophysiology

Because of the similar time dependencies of the behavioral and the neural measure, we wondered if both measures were correlated. We found that the quantities displayed a significant correlation (*r* = 0.33, *p* = 0.0085, *power* = 0.75, Fig. 5C).

## Discussion

We have shown that rhythmic tactile stimulation can significantly improve the recognition of a subsequently presented syllable. The improvement occurred when the syllable started 200 ms after the last vibrotactile pulse. Moreover, we identified an electrophysiological marker of the audiotactile integration, namely the power in the theta frequency band, around 6 Hz, in the right lateral temporal channels and at a delay of 200 ms after the onset of the syllable.

We could furthermore model the time dependency of the syllable recognition score as well as of the neural marker of audiotactile integration through a damped oscillation. Both model fits showed a similar time course, and we found indeed that the behavioral and the neural measure were directly correlated with each other. Because these damped oscillations occurred after the last tactile pulse, they could not result from external stimulation but appear to reflect intrinsic neural oscillations. These results thus helps in disentangling the intrinsic oscillatory activity in the brain from neural activity evoked by rhythmic stimuli such as speech.

Regarding this intrinsic rhythm, our working hypothesis was that the relevant neural activity might be described through an oscillator with a characteristic frequency. Therefore, while we used a tactile stimulation at 5 Hz, any frequency close to the characteristic frequency of the oscillator should be able to entrain the neural activity.

### Syllable Discrimination Score

The syllable discrimination score in the audiotactile conditions was statistically different from that in the audio-only condition at a single audiotactile lag, namely 200 ms. At this delay, the tactile stimulation resulted in a significant improvement in the syllable recognition.

This finding contrasts with previous studies on audiotactile integration regarding syllable recognition, where the effect was found to be restricted to a 50 ms temporal window when the tactile stimulation preceded the audio signal [45]. However, it should be pointed out that this experiment focused on tactile stimuli through a puff of air and how it influenced the perception of syllables with different timing of aspiration (‘ba’ vs ‘pa’). In our experiment, we used vibrotactile pulses that did not carry such information and therefore focused on modulating a rhythm of time windows at which speech processing was more or less effective.

It should also be noted that in related previous papers, the puff of air had an asymmetrical effect on the discrimination between two similar syllables [46, 45]. In our case, we did not observe that the recognition of syllables differed between the different considered syllables, in either of the experimental conditions. Moreover, no condition significantly decreased the syllable discrimination scores. The lack of an adverse effect of the tactile stimulation may aid the application of tactile information in multisensory auditory prosthetics.

Because the delay of 150 ms between the onset of the syllable and the last vibrotactile pulse did not yield a behavioral effect, it appears that the tactile stimulation indeed works in an oscillatory manner, setting up temporal windows where behavior is improved and others where it remains unaffected. A full description of these windows will, in the future, require the consideration of additional audiotactile lags, both shorter and longer than the ones we have investigated here.

### EEG responses

In order to characterize the stimuli we used, we computed the EEG responses to the tactile stimuli as well as to the syllable. The response to the vibrotactile pulses agreed with that found in previous work using the same pulse waveform for somatosensory stimulation, with a strong early response at 48 ms [26]. For repeated vibrotactile events, the EEG response decreased in amplitude, which might be caused by short-term synaptic depression [47]. Similarly to previous work on tactile event-related potentials, we found that the somatosensory-evoked EEG response occurred on the left hemisphere, contralaterally to the location of the right hand to which the tactile stimuli were presented [48].

The EEG response to the syllable presentation exhibited a strong early peak at a delay of 93 ms and a later one at a delay of 207 ms with respect to the onset of the syllable. The topography of the first peak was symmetrical between the left and right hemisphere and suggested an equal contribution from both parts, which is coherent with the asymmetric-sampling-in-time hypothesis [36]. The topography of the second peak was, in contrast, asymmetrical, which is still in line with the asymmetric-sampling-in-time hypothesis, and indicated more right-lateralized activity. Our experiment did not measure speech comprehension per se, but rather syllable discrimination. This task therefore relies on theta rhythm for the slow parsing of acoustic information, which is more prominent in the right auditory cortex [36, 49]. As an alternative explanation for the lateralization of the EEG response, we note that the syllables we were using only differed in their consonants being voiced or unvoiced. The ability to discriminate such voicing cues is typically attributed to the right hemisphere [50].

### Neural marker of audiotactile integration

Using a cluster method, we uncovered a neural marker of audiotactile integration that was also correlated to the behavioral measure. Clustering methods allow for powerful analysis but the interpretation of the obtained results is not straightforward [41]. Indeed, a lot of variability can occur in the boundaries of clusters, making exact inferences about its shape (location, latency and frequency range) unreliable. Therefore, instead of computing all clusters over the whole sets of parameters, we restricted the analysis to certain EEG channels, frequency bands and time windows from certain regions of interest. By doing so, our conclusions do not rely on the exact boundaries of the output cluster but rather on predefined sets established by our working hypotheses. Once a cluster has been established in one of those predefined space-time-frequency region, we reduced the cluster to a single point, thus eliminating cluster boundaries. As a downside of this approach, we had to perform a relatively high number of comparisons to explore several hypotheses, requiring adequate correction for multiple comparisons. As a result, even clusters that exhibited relatively low p-values before correction for multiple comparisons were rejected after multiple-comparison correction and could not be reported in the result. In particular, this was observed for a cluster in the delta frequency range in the right temporal area.

We nonetheless obtained a significant response for audiotactile integration that was located in the right lateral area and contained the power in the theta band. This finding accords with previous work on the theta population in the right auditory cortex, confirming that our task of syllable recognition relied on syllable tracking and that tactile stimulation did indeed affect this process [36]. None of the other hypotheses yielded significant results. In particular, the absence of effects in the delta band is coherent with the roles typically associated with this frequency band, as our syllable-discrimination taks involved no element of syntax or semantics. However, it has been observed that motor delta activity can play a role in speech perception [51]. The absence of an effect in the delta band therefore hints at the fact that either this process does not come into play for syllable discrimination, or that the motor activity in the delta band has not been modulated by our vibrotactile pulses. As for the alpha band, the absence of an audiotactile response indicates that the behavioural result we have observed was not due to an attentional modulation effect, hinting instead at a purely bottom-up processing.

The neural marker of audiotactile integration occurred at a delay of 200 ms after the syllable onset. This delay is in line with our hypothesis that the vibrotactile pulses reset the phase of ongoing theta oscillations, in a similar way as transcranial alternating current stimulation may entrain endogeneous brain oscillations [27]. We also note that only one audiotactile condition, the one with a delay of 200 ms, yielded a significant neural gain. The audiotactile gain in theta activity therefore depended on the alignment between the tactile and the audio signals, as we observed before in related work [26]. This finding might reflect the phenomenon of supra-additivity which may occur when theta oscillations become entrained by the tactile stimulation and are in phase with the syllable [52]. Moreover, it should be noted that no multisensory activity has been detected in the case of the shortest audiotactile lag, thus ruling out the possibility that the observed effect is due to contamination by the somatosensory ERP. In addition, while the GFP of the irregular condition (Fig. 4D) exhibits a later maximum at 400 ms which was not present for the other conditions, it was not statistically significant.

### Damped Oscillation

We were able to describe the time dependency of the behavioral measure and the neural marker of audiotactile integration through damped oscillations. This suggests that the neural activity in the auditory cortex can indeed be partly described as an oscillator, the phase of which is reset through the tactile stimulation. Moreover, both syllable recognition and the neural audiotactile gain exhibited similar temporal dynamics. They both exhibited a phase of about *−*1.16 rad following the tactile stimuli. The first maximum in the syllable recognition score resp. in the audiotactile gain therefore occurred shortly after the last vibrotactile pulse. The largest enhancement of syllable recognition thus occurs if the syllable starts just slightly after the last vibrotactile pulse, in accordance with past work on the subject [26]. The window of enhanced acoustic processing therefore does not coincide exactly with the tactile event. Rather, the vibrotactile pulse may open up a series of windows of enhanced acoustic processing, following a theta oscillation. In addition, the decay times for both the behavioural and the electrophysiological marker were on the same time scale, that of a single period. This evidences that the oscillations dissipate over the course of a few periods.

In addition to the similarity of the obtained model fits, we found that the syllable discrimination scores were indeed correlated to the electrophysiological marker. This reinforces the growing body of evidence that tactile stimuli may modulate speech comprehension through phase reset of endogenous sustained oscillations [27].

## Conclusion

Our previous work on audiotactile speech revealed that comprehension of ongoing speech could be modulated through simultaneous tactile signals linked to the syllable rhythm. The modulation depended in a sinusoidal manner on the time difference between the vibrotactile pulses and the perceptual centers of the syllables [26]. However, it was unclear whether the effect of the tactile stimuli was caused by a reset of ongoing neural oscillations or simply reflected the rhythm set by the successive syllables in the speech stream. Here, we limited the task to syllable discrimination rather than speech comprehension. Because we presented in each trial a single syllable, this could not establish a rhythm as in continuous speech, and we could therefore isolate the effect of rhythm in the tactile stimulation. We demonstrated that establishing a tactile rhythm before presenting a single syllable affected the recognition of that syllable, in a way that could be described by a damped oscillation .A neural marker of the audiotactile integration was found in the theta frequency range, linked to the rate of syllables in continuous speech, and displayed a similar temporal behavior. Together, these results highlight the importance of theta rhythm in syllable perception and are consistent with the hypothesis that tactile signals aid comprehension through resetting the phase of ongoing cortical oscillations in the theta range.

